# A Unified and Interpretable Framework for Evaluating Fluorescence Trace Quality in Transcription Kinetics

**DOI:** 10.64898/2026.01.07.698175

**Authors:** Yiwen Xing, Wei-Tung Lu, Jinhao Liu, Zhiyong Zou, Rui Zhou, Hao Wang, Yusheng Yang, Yabing Yao, Qian Yang, Xiaomin Xu, Hongpeng Zhou

## Abstract

Quantifying transcriptional dynamics from fluorescence traces is a powerful approach to understanding gene regulation, but such analysis critically depends on the quality of the fluorescence signal. Experimental researchers often lack an objective and computationally simple way to assess trace quality before kinetic modeling. In this study, we fill in this gap via systematically investigating two key factors (i.e., signal-to-noise ratio (SNR) and trace length) using synthetic data generated from a composite-state Hidden Markov Model (cpHMM) simulator. By analyzing thousands of simulated traces, we identified quantitative thresholds (SNR ≥ 30 dB and length ≥ 360) beyond which transcriptional dynamics can be reliably captured for kinetic inference. Building on these findings, we further discovered a unified and easily computable quality indicator based on the difference between the first two autocorrelation lags. A threshold value of approximately 0.07 effectively separates reliable from low quality traces, providing a simple yet robust criterion for data selection. Together, these results establish a practical framework for assessing fluorescence trace reliability, offering experimental researchers an interpretable and computationally efficient tool to ensure data quality prior to transcription kinetics modeling.

## 1 Introduction

Transcription kinetics, as a dynamic regulatory process, is crucial not only for normal developmental processes but also in the progression of various diseases [1, 2]. It describes the temporal patterns and rates of RNA synthesis driven by promoter activity, reflecting the stochastic nature of gene expression. Moreover, investigating transcription kinetics allows us to reveal the complex mechanisms underlying gene expression regulation, which has significant implications for cellular reprogramming, gene therapy, and drug design [3, 4, 5]. Investigating these transcriptional dynamics through quantitative modelling and live-cell analysis provides deeper insight into the spatiotemporal interactions between transcription factors and chromatin, which are essential for decoding genome function [6].

The advent of fluorescent protein technology has enabled scientists to fuse fluorescent proteins with target proteins, allowing for the dynamic observation of these fusion proteins in living organisms through fluorescence microscopy [7]. Quantifying transcriptional dynamics through fluorescence traces has become a powerful strategy to uncover gene regulatory mechanisms [8]. Here, fluorescence traces refer to time-resolved measurements of fluorescence intensity at a specific genomic locus in living cells, where the signal arises from nascent RNA transcripts labeled by fluorescent reporter systems (e.g., MS2). By tracking transcriptional bursts in living cells, researchers can infer promoter kinetics, transition probabilities, and transcription initiation rates, offering deep insights into stochastic gene regulation. However, such quantitative inference is highly sensitive to data quality. Experimental fluorescence signals are often affected by different experimental factors and noises. For example, maintaining the brightness and stability of fluorescent proteins in imaging applications remains a significant challenge, as their brightness depends on the molar extinction coefficient and quantum yield, which are influenced by the molecular structure of the proteins and the cellular environment [9]. In fluorescence microscopy experiments, bleaching often impacts the brightness and stability of fluorescent proteins. Over time, the fluorophores undergo bleaching, leading to an exponential decay in fluorescence intensity during the course of the experiment [10, 11]. Furthermore, detector currents and tracking errors are common sources of experimental noise [10]. However, these noises are artefacts in the data and do not represent actual biological processes [10]. These influential factors raise practical concerns about fluorescence trace quality, often leading to unreliable kinetic estimates when data reliability is not properly assessed. Despite its importance, there is still a lack of simple and objective methods to evaluate fluorescence trace reliability before kinetic modeling.

To address this gap, we focused on two fundamental factors that directly influence the reliability of transcriptional fluorescence data—signal-to-noise ratio (SNR) and trace length. SNR provides a natural quantitative measure of noise contamination, while trace length determines whether the recorded signal sufficiently captures the full range of transcriptional dynamics. We generated synthetic fluorescence data from a composite-state Hidden Markov Model (cpHMM) simulator [12]. This simulation framework enables precise control over transcriptional bursting parameters while providing ground-truth kinetic states for benchmarking analytical accuracy. Through extensive simulations and repeated experiments across a wide range of SNRs and lengths, we identified quantitative reliable indicators and threshold values for SNR and trace length separately.

Building upon these findings, we further introduced a unified and easily computable indicator derived from the autocorrelation difference between the first two lag values. This indicator reflects signal smoothness and temporal coherence, offering an interpretable proxy for assessing trace quality without complex modeling. A threshold of approximately 0.07 effectively separates reliable from noisy traces, enabling rapid pre-selection of usable data for downstream kinetic analysis. Rather than proposing a new kinetic inference method, our goal is to systematically characterize the operating regime in which widely used autocorrelation-based analyses remain reliable. Together, this work establishes a quantitative and interpretable framework for evaluating fluorescence trace reliability, bridging the gap between data generation and trustworthy transcriptional modeling.

## 2 Methods

In this section, we outline the methodology used for generating synthetic fluorescence intensity data and performing autocorrelation analysis to estimate polymerase dwell time.

### 2.1 Synthetic Fluorescence Traces Generation

A simulator based on a Hidden Markov Model (HMM) was utilized to generate synthetic fluorescence intensity traces. The methodology for generating synthetic data was presented by Hoppe et al. (2020) [13]. Specifically, the data generation process can be divided into two steps. First, synthetic promoter sequences are generated using a Markov process. We produce traces of predetermined length based on a known transition matrix, thereby constructing synthetic promoter sequences. Figure 1(a) illustrates the temporal dynamics of a sample promoter trace (ON/OFF), which forms the fundamental basis for the generation of synthetic fluorescence signals. Second, synthetic fluorescence traces are generated from the simulated promoter states. Following Hoppe et al. (2020) [13], a pretrained HMM model using promoter sequences as input can accurately capture mRNA transcription activity and generate realistic fluorescence signals. As shown in Figure 1(b), the synthetic fluorescence trajectories correspond to the promoter activity depicted in Figure 1a. In the current implementation, the noise level is set to 0, ensuring that the generated fluorescence traces as the baseline.

**Figure 1.**
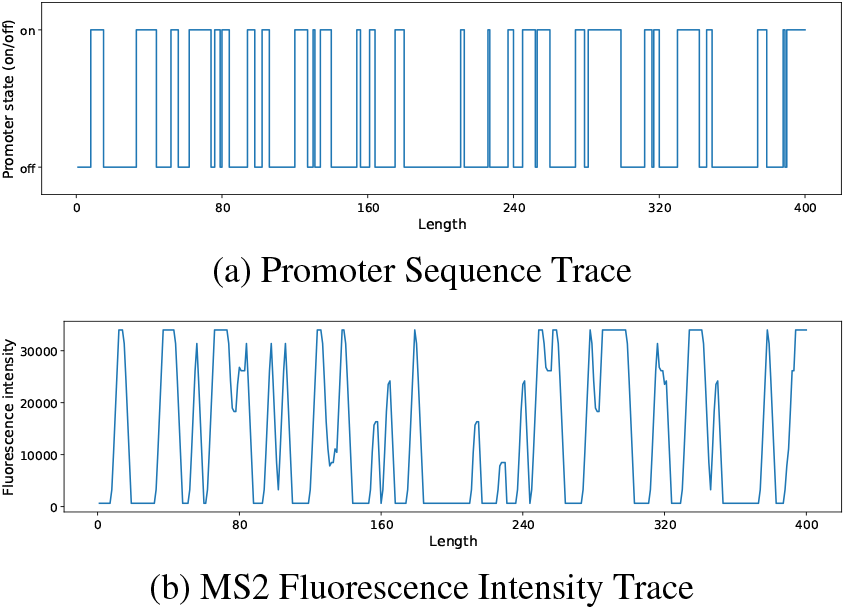
A snapshot of Synthetic data.

The noisy synthetic traces are generated with different signal-to-noise ratio (SNR). SNR is a measure used to quantify the level of a desired signal relative to the level of background noise. A high SNR indicates that the useful information in the signal far exceeds the noise, while a low SNR indicates that noise occupies a larger proportion of the signal. It is commonly expressed in decibels (dB) and is defined as:

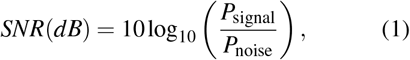

where *P*_signal_ is the power of the signal, which can be calculated as the mean squared value of the fluorescence intensity signal:

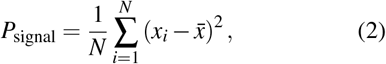

Here, *x*_*i*_ represents the individual data points of the signal, and 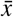 is the mean of these data points. N is the total number of data points in the signal. *P*_noise_ is the power of the noise, calculated similarly as the mean squared value of the noise component added to the signal:

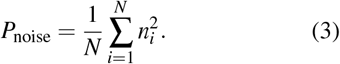

where *n*_*i*_ represents the individual noise data. By adjusting the noise power *P*_noise_, different SNR levels can be simulated, allowing us to investigate how varying levels of noise impact the accuracy and reliability of fluorescence intensity measurements. In this paper, noise-free fluorescence intensity signals (*SNR* = ∞) were generated as the baseline. Then, different levels of noise were added using a Gaussian distribution with SNR ranging from 0dB to 60dB to simulate realworld data variability. As illustrated in Figures 2, the fluorescence intensity traces show how increasing levels of noise affect the signal.

**Figure 2.**
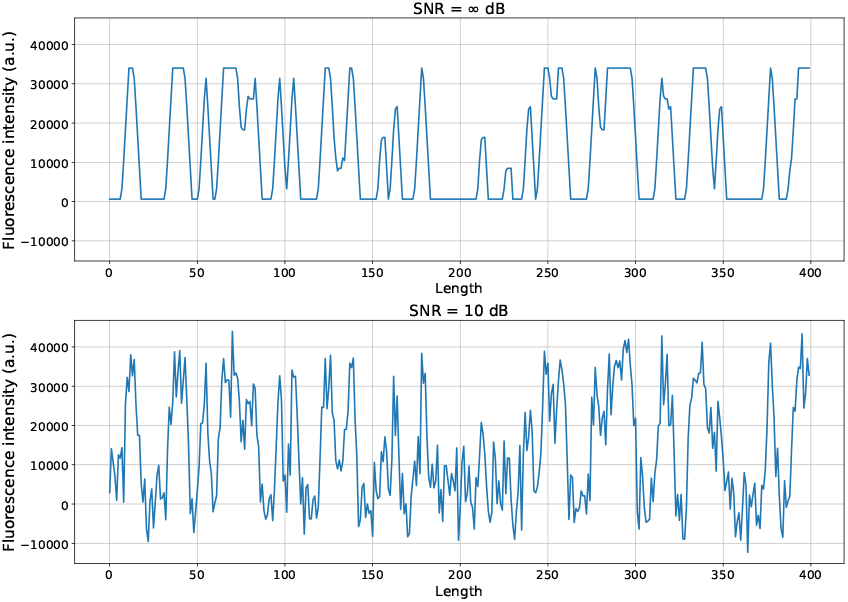
Synthetic fluorescence signals with different noise

### 2.2 Autocorrelation Analysis for Polymerase Dwell Time Estimation

Dwell time is an indicator of how long the polymerase remains engaged with the DNA template at each step before proceeding to the next, thereby reflecting the overall dynamics of the transcription process [12, 14].

Autocorrelation of a signal provides a measure of how a signal at one point in time is related to the same signal at a different time. By calculating the autocorrelation function R(τ), where τ represents the lag or time delay, we can assess how the fluorescence signal correlates with itself over time. The autocorrelation function is defined as:

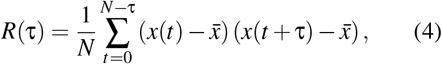

where *x*(*t*) is the fluorescence signal, 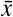 its mean, and *N* the length of the signal.

In transcription, the autocorrelation function is useful because it reflects the processes of RNA synthesis initiation, elongation, and termination during transcription [14]. The dwell time can be inferred from the autocorrelation function as it often corresponds to the characteristic time where the autocorrelation begins to decay significantly.

As shown in Figure 3, the autocorrelation values shows a rapid initial decay followed by stabilisation. This pattern is indicative of the underlying system dynamics, where the initial decrease in autocorrelation reflects the decay of short-term correlations, and the subsequent plateau represents stable long-term correlation. The indicated “Dwell Time” on the curves represents the characteristic time scale at which this stabilisation occurs.

**Figure 3.**
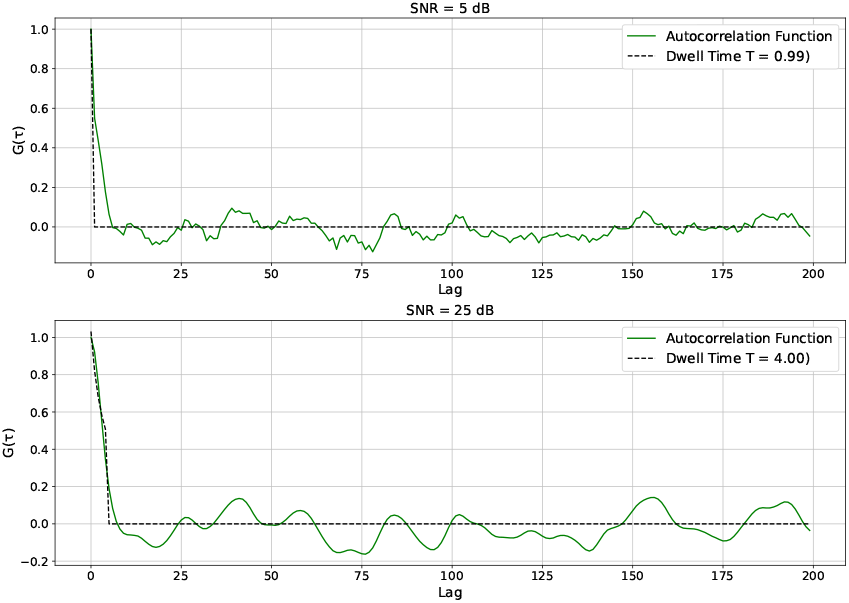
Dwell time prediction by fitting autocorrelation values.

To accurately determine the dwell time for each curve, we use a parametric fitting function based on Larson et al. (2011) [14]. Here, τ denotes the discrete *lag* (time delay) measured in multiples of the sampling interval Δ*T*; thus *R*(τ) is the empirical autocorrelation of the fluorescence signal at lag τ. We approximate *R*(τ) with the model *G*(τ) and estimate *T* by fitting *G* to *R* over τ ∈ [0, τ_max_] via nonlinear least squares:

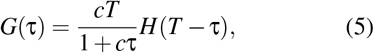

where *H*(*x*) is the Heaviside function, defined as:

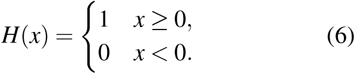

The goal of curve fitting is to find a set of parameters c and T such that the model function G(τ) best matches the calculated autocorrelation values R(τ). To achieve this, we use nonlinear least squares to minimise the sum of squared differences between the two functions:

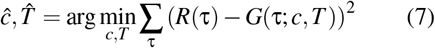

Through this method, we can estimate the dwell time T of the signal. It is important to emphasise that the dwell time T obtained using this method is not a specific point on the autocorrelation curve, such as a turning point, but rather a parameter that defines the shape of the fitted curve. As shown in Figure 3, the value of T is displayed in the legend as part of the fitted curve description. This parameter T governs the transition of the autocorrelation function from its initial rapid decay to stabilisation.

## 3 Experimental Design

In this section, we elaborate on the experimental design, including the setup of the SNR and trace length experiments, as well as the rationale behind discovering the reliability indicator.

### 3.1 SNR Dependence Analysis

In the study of transcription kinetics, SNR is a crucial metric for assessing signal quality. A high SNR ensures that the data we obtain are closer to the ideal scenario, reducing the impact of noise on data interpretation. Therefore, determining a reasonable SNR range can enhance the credibility and accuracy of the data, thereby improving the precision of the research results. In the SNR experiment, we compare synthetic fluorescence intensity data with a SNR range of 0-60dB to determine the SNR level required to make the data interpretation similar to that of noiseless (Ground Truth) data under ideal conditions.

To better understand how various SNR levels influence dwell time, we present the distribution of dwell times for 800 curves at each SNR level using box plots. Autocorrelation traces offer several advantages: they not only provide a clear visual representation of the median, quartiles, and overall distribution of the data, offering a comprehensive overview of the data’s spread, but they also reveal the variability within the data. Additionally, box plots effectively identify outliers, enabling us to detect dwell time conditions that deviate from the Ground Truth. While box plots do not directly display the standard deviation, they convey the interquartile range (IQR), which allows for an estimation of the standard deviation and, consequently, the data’s dispersion. Thus, box plots serve as a robust tool for comprehensively analysing the impact of different SNR levels on dwell time (see details from Figure 5 and Figure 7 in Section 4).

### 3.2 Trace Length Dependence Analysis

A sufficient number of sampling points is essential to accurately capture transcriptional dynamics. The purpose of finding the shortest required trace length is to help scientists determine the appropriate sampling trace length during data collection. This ensures that the complete characteristics of the gene transcription curve are preserved, thereby improving data interpretability and research accuracy. By determining this critical trace length, not only can experimental resources be conserved, but unnecessary data redundancy can also be avoided, making the experimental design more efficient and precise.

As shown in Figure 4, it can be observed that the predicted dwell time also varies with trace length. Autocorrelation fitting serves as an effective approach to identify such an appropriate indicator for evaluating trace length. When the trace length is too short, the autocorrelation function fails to accurately capture the dwell time, leading to large variations and unstable estimates.

**Figure 4.**
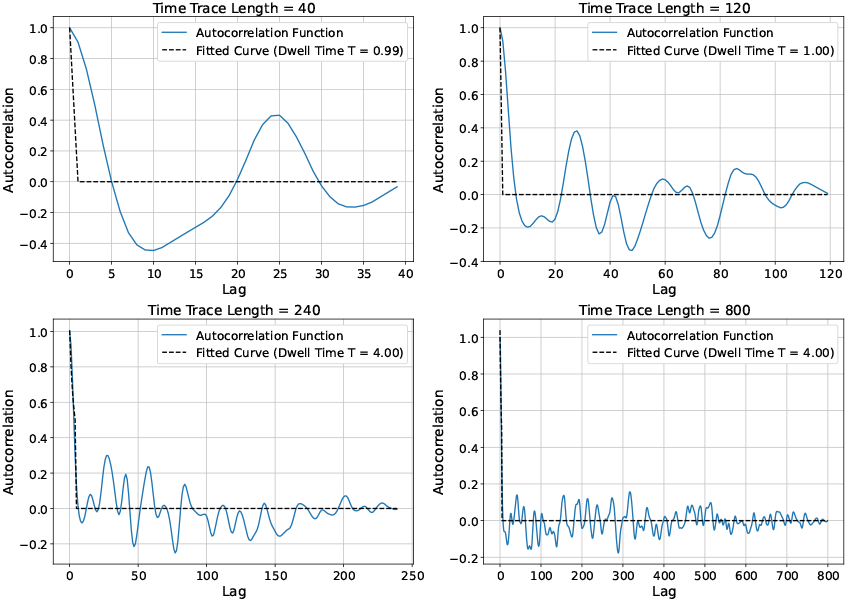
Autocorrelation fitting at different trace lengths.

**Figure 5.**
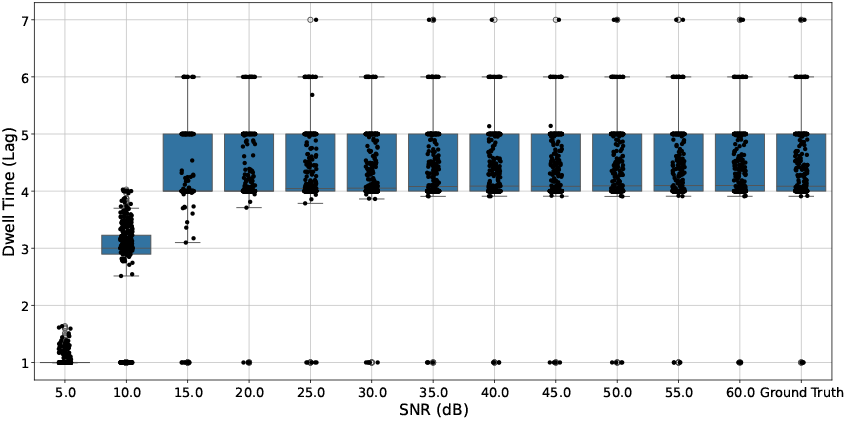
Dwell time distribution across different SNR levels.

In this experiment, we evaluated data across various trace lengths ranging from 40 to 800 points to determine the minimum duration required for obtaining interpretable and reliable results. The convergence of the predicted dwell time was assessed by identifying the point at which the estimates became stable, indicating that further increases in trace length no longer produced significant changes. This stabilization reflects that the data have reached a level of reliability and consistency comparable to the ground truth, thereby defining a critical duration for data acquisition. To determine the appropriate trace length, we employed a two-step approach: first, dwell times were calculated for each curve across different trace lengths and visualized as box plots to observe distributional patterns; second, the mean dwell time values were analyzed by computing the relative rate of change between consecutive lengths. A relative change rate below 2% was considered indicative of convergence and stability.

### 3.3 Discovery of a Unified Reliability Indicator

In transcription dynamics experiments, when using the autocorrelation function to learn the kinetic dynamics [14, 15], the focus is typically on how autocorrelation values change at different lags. However, for biologists, controlling the observation trace length is relatively straightforward, whereas measuring SNR is quite challenging. Although the appropriate range of SNR can be determined from previous experimental results of section 3.1, measuring the true SNR in real situations is often difficult. Therefore, we aim to provide a more direct indicator to indirectly assess data reliability, making it easier for biologists to evaluate the stability of their data.

Our observations indicate that most autocorrelation curves exhibit a noticeable downward trend before reaching the dwell time. Specifically, autocorrelation values decrease with increasing lag until the dwell time is reached. To quantify this behaviour and assess data reliability, we propose a new metric based on the difference between different lags. This metric focuses on the difference between Lag=0 and Lag=1 as an indicator of data reliability.

We have three reasons for choosing this indicator. First, At Lag=0, the autocorrelation value is always 1, representing the correlation of the signal with itself at the same time point. This value is inherently maximal, meaning that any deviation from this point will be negative. Second, as shown in Figure 4, the slope between Lag=0 and Lag=1 is the steepest among all adjacent lags. This indicates that the rate of change in autocorrelation values is most pronounced between these two lags, making it easier to distinguish different scenarios. Hence, Lag=1 provides a sensitive measure for detecting variations in data reliability. Thirdly, Using lags greater than 1 may introduce more significant errors, especially in cases of high signal noise or insufficient observation time. Larger lags can lead to greater fluctuations in the autocorrelation curve, thereby increasing measurement errors. By focusing on Lag=1, we can reduce such errors and enhance the accuracy of our analysis.

We have four specific calculation procedures: *(1) Calculate the Autocorrelation Function:* Initially, compute the autocorrelation function based on the fluorescence signal data to obtain autocorrelation values for each lag. This step reveals the correlation of the signal over different time points and provides the necessary data for further analysis. *(2) Calculate the Mean Autocorrelation Curve:* For various SNR levels or trace lengths, compute the average of all autocorrelation curves. This averaging process helps smooth out noise and provides a clearer reflection of the overall trend. *(3) Calculate Differences Between Lags:* Focus particularly on the difference between Lag=0 and Lag=1. This difference serves as a key indicator for evaluating data reliability, as it represents the most significant change in autocorrelation values. *(4) Observe Changes:* Plot a line graph to analyse the variation in the differences of the autocorrelation function.

## 4 Results and Discussion

In this section, we present and discuss the results of experiments designed to evaluate the effects of signal-to-noise ratio (SNR) and trace length on the reliability of synthetic fluorescence data. We first analyze how dwell time distributions vary across different SNR levels and determine the minimum SNR required to ensure high-quality traces. We then examine the influence of trace length on dwell time estimation and identify the minimum length needed for stable measurements. Finally, we proposed a unified and reliable indicator which can work under different experimental conditions. The code is available at https://github.com/DianaLu-2022/GeneKineticsReliability/tree/master.

### 4.1 SNR Experiment Results

SNR is a critical factor influencing the quality of gene transcription data. To analyse how dwell time changes under different SNR levels, we conducted experiments and visualised the distribution of dwell times across various SNR levels using box plots.

#### 4.1.1 Variation in Dwell Time Distribution across SNR Levels

The experimental results indicate that SNR levels significantly affect the distribution of dwell times (as shown in Figure 5). Under low SNR conditions (e.g., 0 dB), the distribution of dwell times is broad, with high variability, particularly between 1 and 2 time points (lag), and a notable presence of outliers. As the SNR increases (e.g., 5 dB, 10 dB), the distribution range expands, and the median shifts upward, demonstrating the significant impact of noise on the transcription process. When the SNR reaches 15 dB or higher, the distribution of dwell times begins to stabilise, concentrating between 4 and 6 time points, with a noticeable reduction in outliers. This indicates a significant improvement in data reliability. At high SNR levels of 30 dB and above, the dwell time distribution becomes consistent, closely resembling the Ground Truth, suggesting that the impact of noise has been effectively minimised.

#### 4.1.2 Identification of the SNR Stability Threshold

As shown in Figure 6, by comparing predicted dwell time of across traces with different SNR levels, the stable point was determined where the mean dwell time changes by less than 0.02 (i.e., 2%) between successive lengths, which occurs at 30 dB. It can also be found that at SNR level of 30 dB and above, the predicted dwell time is close to the Ground Truth. This indicates that at these SNR levels, the data reliability is highest and is suitable for transcription analysis. These findings regarding SNR levels and dwell time can serve as benchmarks for evaluating signal quality, providing a reliable metric to assess whether the data is sufficiently robust for accurate transcription analysis.

**Figure 6.**
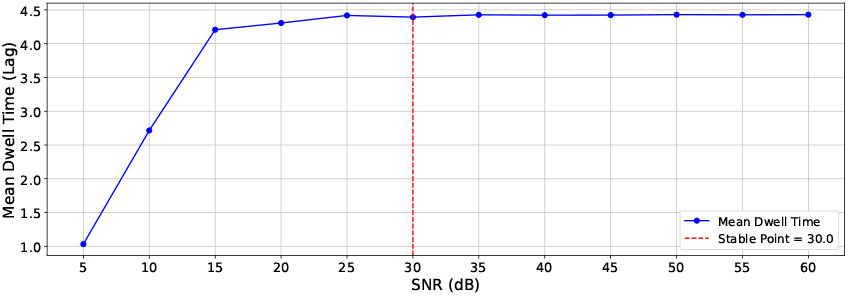
Stabilisation of mean dwell time at increasing noise.

**Figure 7.**
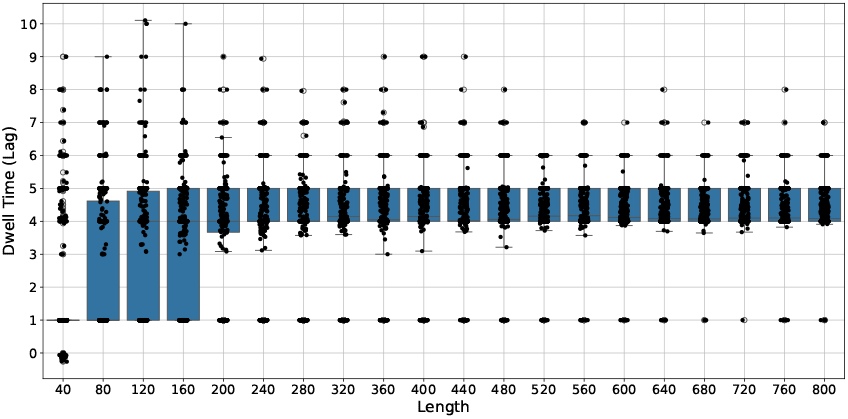
Dwell time distribution across different trace lengths.

### 4.2 Trace Length Experiment Results

In gene transcription analysis, the length of the Trace is a crucial factor that affects the quality and interpretability of the data. To understand the trend of gene transcription dwell time across different trace lengths, we conducted multiple experiments and visualised the distribution of dwell times using box plots and mean value trend graphs.

#### 4.2.1 Distributional Trends across Different Trace Lengths

As shown in Figure 7, the distribution of dwell times becomes more stable as the time trace length increases. For shorter trace lengths (60 to 160 points), dwell time variability is greater, with a broader distribution and more outliers. This indicates that data stability is lower, and dwell time measurements are less reliable at shorter trace lengths. Specifically, when the trace length is smaller 160, the dwell time of the repeated experiments has a larger variance with several outliers, indicating significant random fluctuations. However, as the trace length increases to 200 points and beyond, the distribution of dwell times begins to converge with a noticeable reduction in outliers.

#### 4.2.2 Determination of the Minimum Stable Trace Length

By analysing the relationship between mean dwell time and trace length, we identified 360 time points as the threshold at which dwell time begins to stabilise. As shown in Figure 8, the stable point was determined where the mean dwell time changes by less than 0.02 (i.e., 2%) between successive lengths, which occurs at 360 points. This point marks a critical benchmark for evaluating the quality of experimental data. This suggests that a trace with length beyond 360 can further enhance data stability and reliability. Therefore, selecting an appropriate trace length, ideally greater than 360 points, is crucial in ensuring data quality and reliability in practical experimental design.

**Figure 8.**
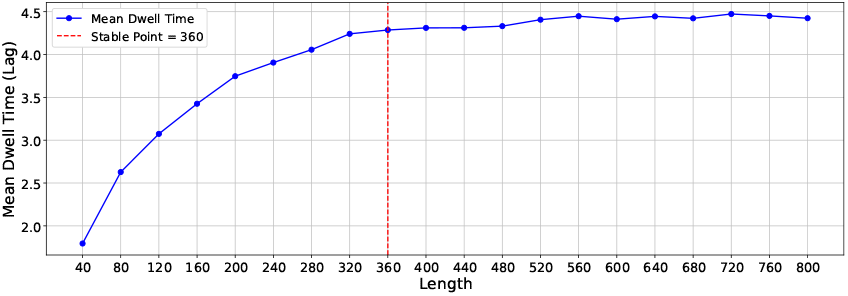
Stabilisation of mean dwell time at increasing trace lengths.

### 4.3 Identification of a Unified Reliability Indicator

Following the proposed lag difference indicator as described in Section 3.3, we conducted a detailed experimental analysis. This new metric aims to simplify the process of assessing data reliability, making it more user-friendly for biologists. Through our experiments, we systematically assessed the performance of this indicator under varying SNR levels and different trace lengths to verify its accuracy and stability.

For the SNR experiment, as shown in Figure9, the difference between Lag=0 and Lag=1 decreases significantly as the SNR increasing. At the stable point of 30 dB, identified in Section 4.1.2, the autocorrelation value difference ireaches approximately 0.0707, which can serve as a new evaluation metric for data reliability. Notably, when the SNR reaches 30 dB or higher, this difference closely approaches the value observed in the ground truth, indicating that at high SNR levels, noise influence is substantially reduced and the fluorescence traces exhibit higher reliability.

For the trace length experiment, as shown in Figure 10, the autocorrelation difference at the stable point (length = 360) is 0.0702, which closely matches the value obtained from the SNR experiment in Figure 9. Therefore, a threshold value of approximately 0.07 can be established as a unified reliability indicator for evaluating trace quality. By applying this metric, researchers can rapidly and objectively determine whether their fluorescence traces meet reliability standards, particularly in experiments affected by varying noise levels.

**Figure 9.**
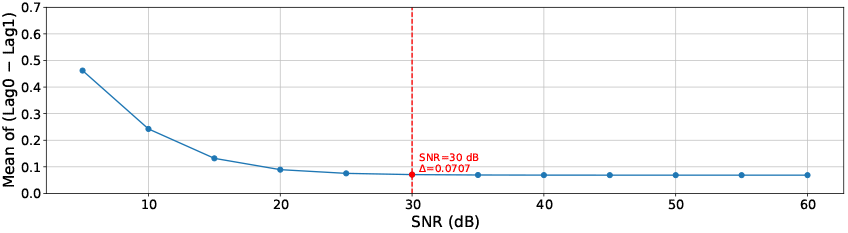
Difference of autocorrelation value between Lag 0 and lag 1 at various SNR levels.

**Figure 10.**
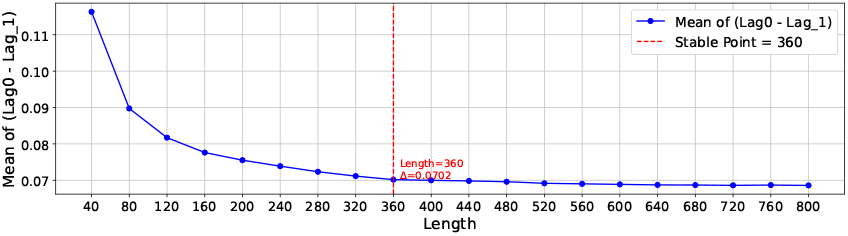
Difference of autocorrelation value between Lag 0 and lag 1 at various trace lengths.

For clarity, this study focuses on autocorrelation-based analysis, a simple, interpretable, and widely used approach in transcription imaging studies. The results should therefore be interpreted in the context of applications that rely on autocorrelation-based methods. In addition, we have implemented our approach in real-data evaluations and applied it to sequence selection, as reported in [16].

## 5 Conclusion

This study introduces a simple and reliable indicator to assist experimental researchers in evaluating the quality of transcriptional fluorescence traces. Through systematic simulations, we identified that traces with a signal-to-noise ratio (SNR) of at least 30 dB and a minimum length of 360 time points can robustly capture transcriptional dynamics consistent with the ground truth. We further propose an intuitive quality metric based on the autocorrelation difference between Lag = 0 and Lag = 1, with a threshold of approximately 0.07 distinguishing reliable traces from noisy ones. This approach eliminates the need for complex modelling or manual inspection, providing a computationally efficient and interpretable framework for large-scale data screening. While currently demonstrated on simulated data, this indicator offers a promising direction for real-time trace quality control in experimental transcription imaging.

